# Bias in group-level EEG microstate analysis

**DOI:** 10.1101/2022.11.07.515464

**Authors:** M Murphy, J Wang, C Jiang, L Wang, N Kozhemiako, Y Wang, the GRINS consortium, JQ Pan, SM Purcell

## Abstract

Microstate analysis is a promising technique for analyzing high-density electroencephalographic data, but there are multiple questions about methodological best practices. Between and within individuals, microstates can differ both in terms of characteristic topographies and temporal dynamics, which leads to analytic challenges as the measurement of microstate dynamics is dependent on assumptions about their topographies. Here we focus on the analysis of group differences, using simulations seeded on real data from healthy control subjects to compare approaches that derive separate sets of maps within subgroups versus a single set of maps applied uniformly to the entire dataset. In the absence of true group differences in either microstate maps or temporal metrics, we found that using separate subgroup maps resulted in substantially inflated type I error rates. On the other hand, when groups truly differed in their microstate maps, analyses based on a single set of maps confounded topographic effects with differences in other derived metrics. We propose an approach to alleviate both classes of bias, based on a paired analysis of all subgroup maps. We illustrate the qualitative and quantitative impact of these issues in real data by comparing waking versus non-rapid eye movement sleep microstates. Overall, our results suggest that even subtle chance differences in microstate topography can have profound effects on derived microstate metrics and that future studies using microstate analysis should take steps to mitigate this large source of error.

## Background

Multichannel electroencephalographic (EEG) data can be modeled as a set of semi-stable recurring voltage topographies called microstates. Each microstate persists for 50-150 milliseconds before transitioning to a different microstate. Multiple studies have demonstrated that only a small number of unique microstates are needed to explain most of the variance in the EEG signal^1^. These microstates are extracted via data-driven clustering, most commonly modified *k*-means^2^ or atomize and agglomerate hierarchical clustering^3^. While different studies have reported different sets of microstates, four canonical microstates (A, B, C and D) have been widely replicated in studies of resting state EEG^4^. The neurobiological generators of these microstates remain unclear, but evidence suggests that each microstate corresponds to a distinct large-scale neural assembly that overlaps with, but is distinct from, a resting state network as measured by functional magnetic resonance imaging^5^.

Microstate analysis has been used to study normal human physiology as well as psychiatric and neurological disease^6–9^. However, comparing results across studies is often difficult because of multiple analytic decision points during both the extraction of microstate topographies and when “backfitting” these topographies to the original signal data to derive various microstate parameters, as described below. Critically, derived microstate parameters are necessarily dependent upon the modelling and measurement of microstate topographies. Despite these challenges, there are results which appear to be consistent across studies – for example, that schizophrenia may be associated with increased canonical microstate C^7,10,11^.

In many studies using microstate analysis to identify differences between groups, microstate topographies are first extracted from each group independently (see for example ^6–8^). Then, authors compare the sets of topographies and align them based on spatial correlation and similarity to canonical microstate topography. Each group is then backfit with its own topographies. Parameters such as microstate duration, occurrence, explained variance, and transition properties are calculated from these fits and then compared across groups. Conceptually, groups may vary in their characteristic maps (if fundamentally equivalent underlying states manifest with altered scalp EEG topographies), in any of the derived metrics (for example, if one microstate occurs more frequently) or in both domains.

Microstate topographies calculated from different data sets may be broadly similar but are typically not identical. That is, canonical microstate A derived from one data set will not be identical to canonical microstate A derived from a different data set. Multiple studies have shown the canonical microstates C and D have a more varied topography both within and between studies than do microstates A and B^5,10,12,13^. It is unclear to what extent subtle differences in microstate topography may impact subsequently estimated parameters.

When considering group differences, an alternative approach is to derive a common set of microstates (typically but not necessarily from all individuals in a study) and use that common set for backfitting and calculation of microstate parameters. We speculated that the former approach may inappropriately capitalize on chance group differences in microstate topologies, especially when applied to relatively small samples, and thus lead to spurious differences in derived parameters which no longer reflect “apples-to-apples” comparisons. In contrast, using a common set of microstates should reduce the risk of reporting spurious group differences (referred to below as “combined map” versus “subgroup maps” conditions).

We tested eyes-closed, resting state EEG data from 30 healthy control participants. For each of 5,000 replicates, we randomly divided this group into two equal-sized subgroups. Using the Luna analysis package (developed by S.M.P.), we initially calculated microstate parameters by either 1) fitting all the data to a single set of microstates derived from the pooled (*N*=30) data (“common map”), or 2) fitting the data from each (*N*=15) subgroup to a set of microstates derived only from that subgroup (“subgroup maps”). For the subgroup-derived microstates, in keeping with previous work, we paired each microstate with the microstate in the other subgroup it most closely resembled (see Methods). We then calculated whether there were statistically significant (*p* < 0.05) differences in any of five microstate parameters (coverage, duration, occurrence, variance explained, mean spatial correlation) between the two subgroups.

Below we arbitrarily refer to the randomized groups as “cases” and “controls”, for convenience and to mimic a typical analytic context, although as noted above, all original data are from healthy controls. As group labels are assigned under the null hypothesis of no group differences, only 5% of replicates are expected to be significant at *p* < 0.05, if the procedure appropriately controls false positive rates. Distinct from testing for group differences in microstate parameters, we also tested for case/control differences in the microstate topographies themselves, based on summaries of pairwise distances between 30 individual-level maps, e.g. asking whether case/control pairs had more dissimilar maps than case/case or control/control pairs on average (see Methods). Figure 1 shows an overview of the simulation and testing procedure, repeated 5,000 times.

**Figure 1.**
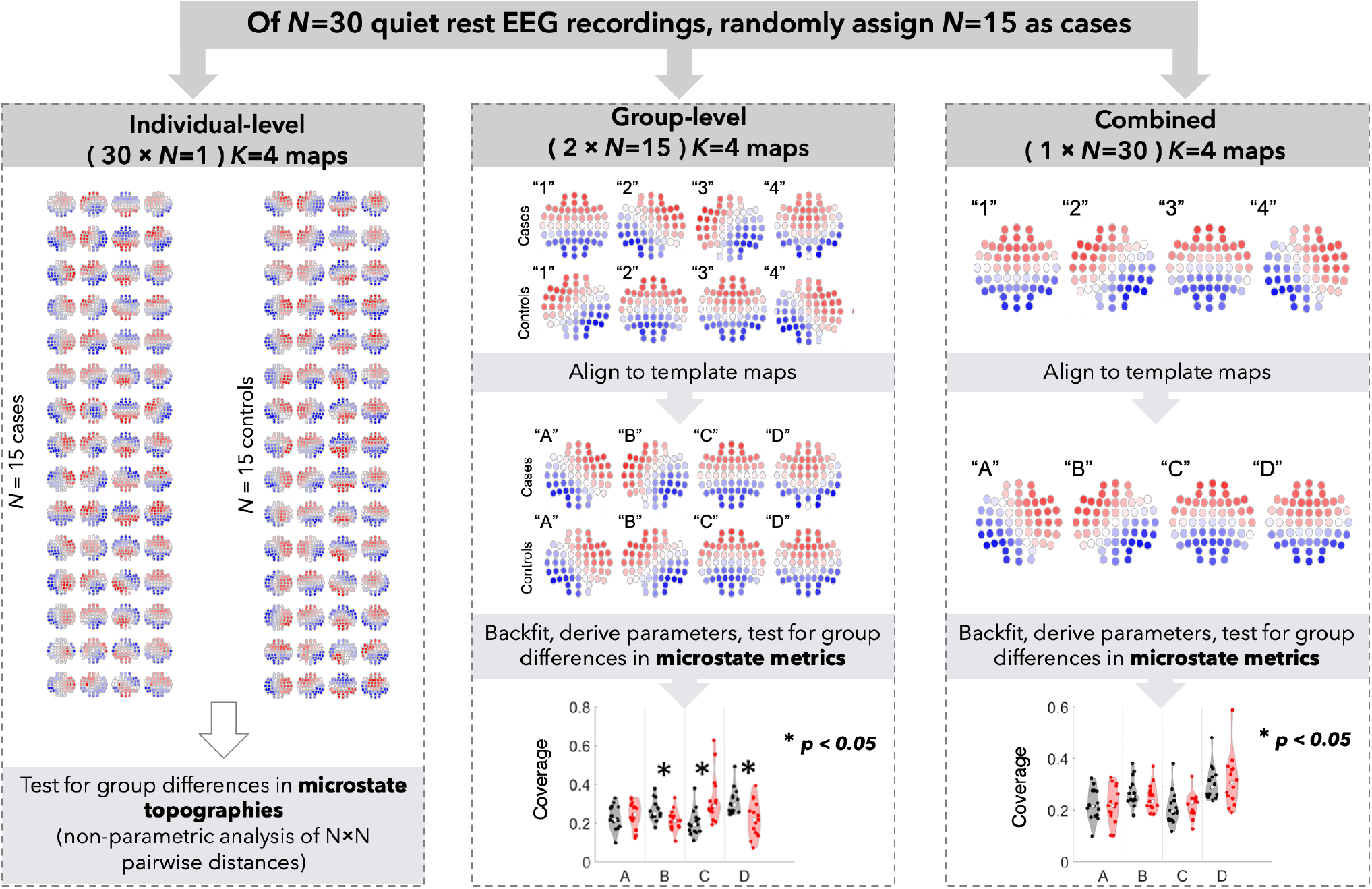
Schematic of the study. Red violin plots are “cases” and black plots are “controls”.

## Results

### Using subgroup specific maps leads to inflated type I error rates

By design, the primary simulation procedure did not induce systematic differences between the topographies of randomly assigned cases and controls. Consistent with this, for all tests comparing the topographical similarity of case and control maps (based on individual-level, 30 × *N*=1 segmentation) we observed type I error rates close to their expected value, i.e. 5% (data not shown). Further, based on subgroup (2 × *N*=15) maps, cases and controls typically showed very high spatial correlations with respect to i) to the assigned canonical template map (median *r* = 0.98), ii) to the matched combined (1 × *N*=30) map (median *r* = 0.99), as well as iii) to each other (median *r* = 0.95). That is, as expected, randomly assigned cases and controls typically exhibited similar *K* = 4 microstate topographies. Further, the median correlation of microstates between subgroups were 0.90, 0.93, 0.95, and 0.97 for canonical microstates A-D respectively, which is comparable to values reported in ^6–8^.

In contrast, despite similar topographies, error rates for group differences in derived microstate metrics were markedly inflated when using subgroup maps, but not when using a common map (Figure 2a,b). For the shared microstate analyses, 5.01% of all comparisons were statistically significant at a *p* < 0.05 threshold and the median number of false positives per permutation was 0 (of 20 tests performed). In contrast, when using subgroup maps, 31.11% of all comparisons were statistically significant at a *p* < 0.05 threshold and the median number of false positives per permutation was 6 of 20. Based on a conservative Bonferroni-corrected error rate of 0.05 / 20, the experiment-wide error rate was 68.78% when using subgroup maps, versus 3.72% when using a combined map. This bias specifically resulted from using different maps between groups: similar to using a combined map, if using the same set of subgroup maps for all individuals (for example, only case-derived maps when backfitting both cases and controls), type I error rates were controlled at their expected values (data not shown).

**Figure 2.**
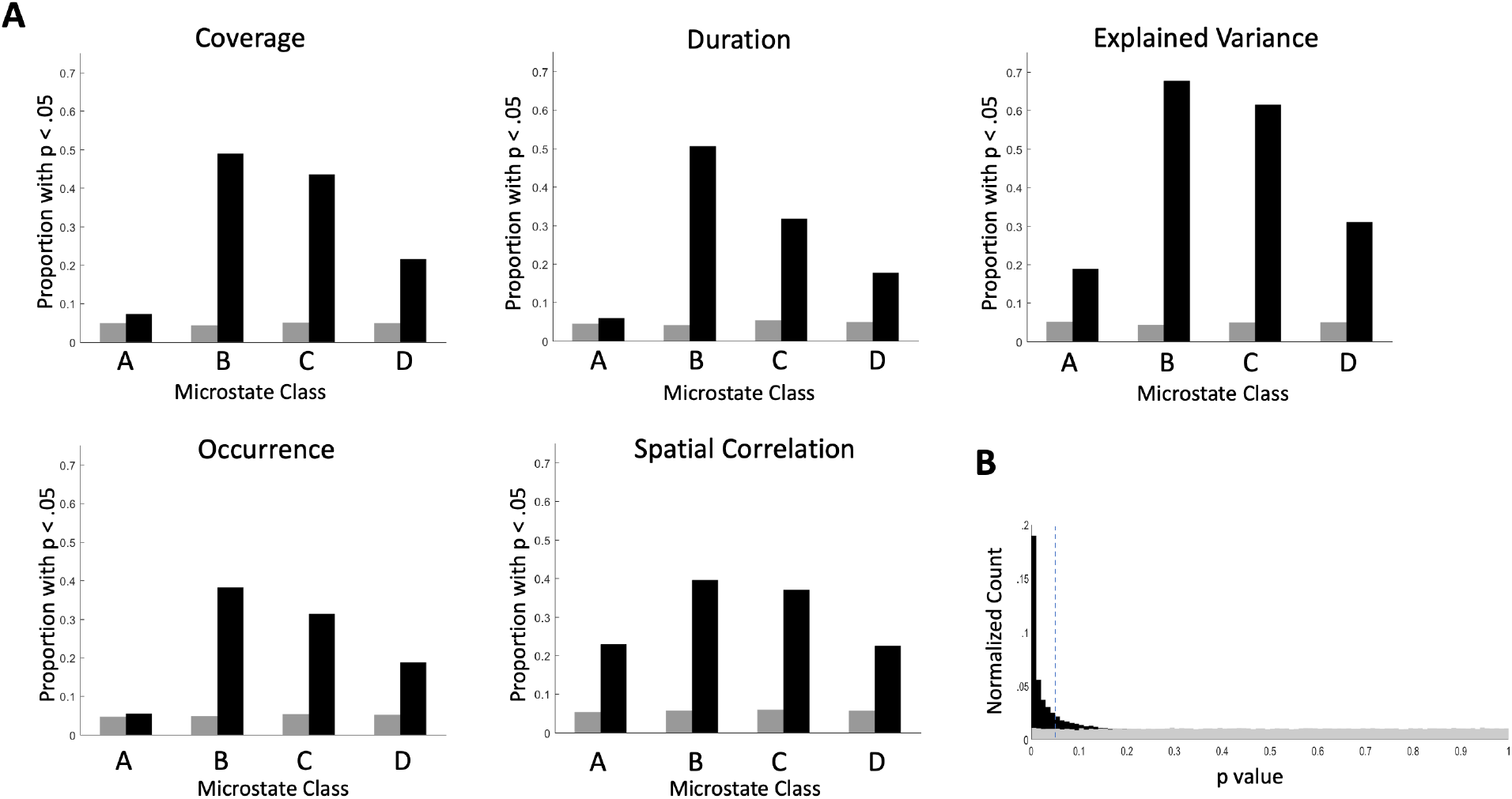
Subgroup maps show consistently higher type I error rates than combined maps. **A)** Proportion of permutations with false positives for each microstate and parameter. Gray bars are combined maps and black bars are subgroup derived maps. **B)** Histogram of p-values for all comparisons in all permutations. The dotted line indicates p = 0.05. There is a large peak of p values less than 0.05 In the subgroup derived maps.

Investigators have long been aware that differences between microstate topographies in different subgroups complicate the interpretation of subgroup differences in microstate parameters^10,12,14,15^. One widespread approach to this complication is to measure the spatial correlation of the microstate maps between subgroups. These correlations are often quite high (for example, see da Cruz et al.^7^, where all correlations are > 0.91). These high correlations are then used to explicitly justify using subgroup-specific templates. We note that we observed elevated type I error rates despite spatial correlations that were comparable with values reported by other groups. Furthermore, we restricted our analyses to the 1,026 replicates with the closest case-control topographies, defined as i) case-control spatial correlations *r* > 0.95 for all four states, ii) no ‘suboptimal’ template map assignments (see Methods) and iii) no significant (*p* < 0.01) case-control differences for tests of topographical similarity. Even in this set, the average error rate was still inflated for analyses based on subgroup maps (16.97%) and but not shared maps (4.99%). Of note, error rates for microstate C were particularly inflated for all five metrics (44.6%) in this reduced set. Taken together, these results suggest that the widespread practice of using spatial correlations to support the use of subgroup derived maps can still be prone to inflated rates of type I error even when subgroup maps are highly correlated.

Overall, inflated error rates were most apparent for comparisons involving microstate B and C, and to a lesser extent D. That microstate A had lower rates of type I error (at least for some metrics) likely reflects different levels of sample-specific variability in topographies, peculiar to this particular set of recordings and choice of *K*=4, rather than general properties of microstate analysis or neurophysiology.

As a concrete example, Figure 3 shows the maps and coverage metrics obtained from one simulation replicate. Although maps are visually similar and have high spatial correlations both between groups and with the template maps, they are not identical, as the plot of map differences shows (Figure 3a). Consequently, although most coverage parameters derived from subgroup and combined maps are correlated with each other, they can also diverge (Figure 3b). Further, this divergence occurs differentially between case and control groups, leading to a spurious difference in coverage for microstate B, C, and D (Figure 3c).

**Figure 3.**
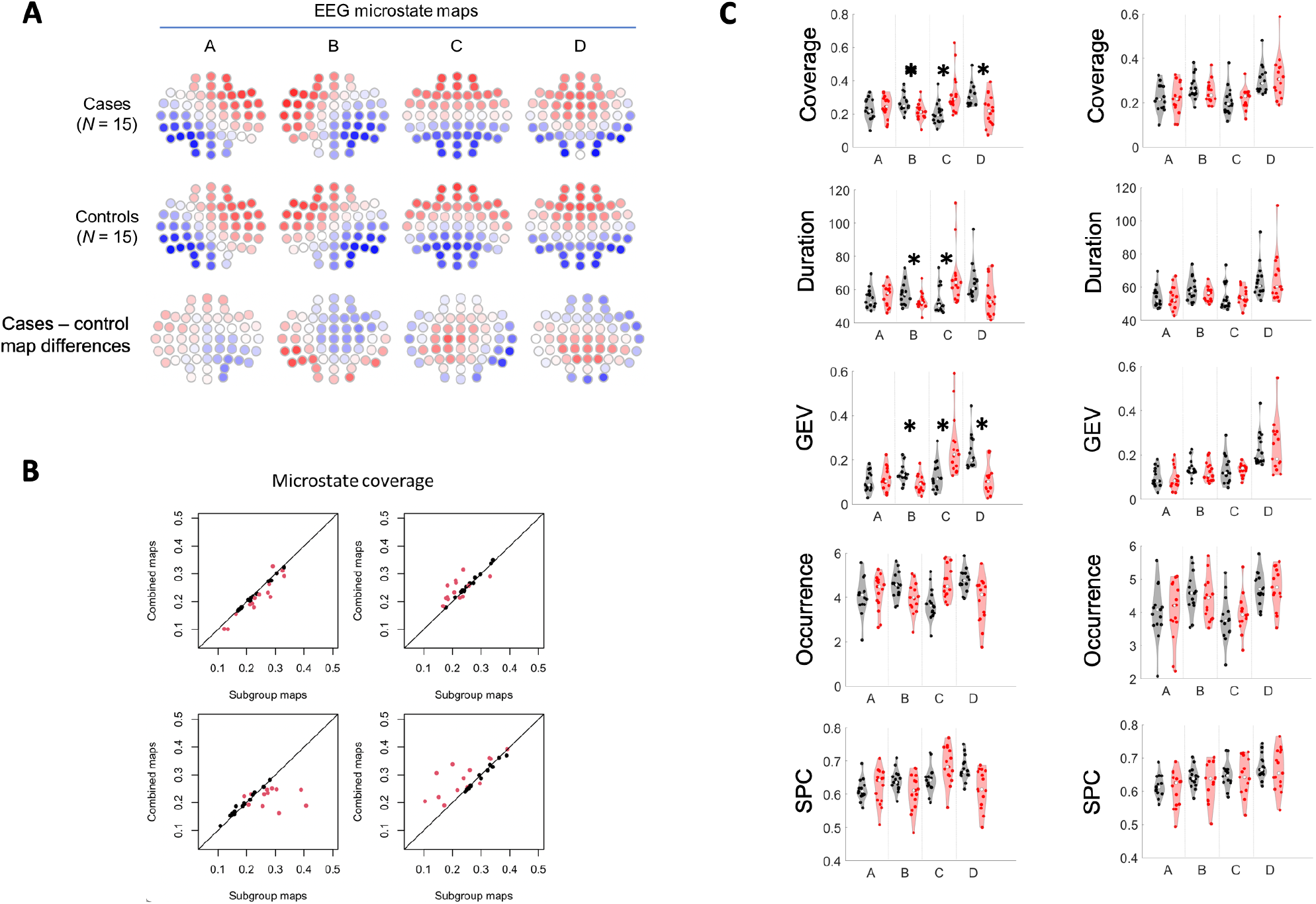
An example of elevated type I errors when using subgroup maps for a single permutation. **A)** Subgroup derived case and control maps and the difference between them. **B)** Correlations between the coverage metrics calculated with subgroup maps for cases (red) and controls (black) for each microstate. **C)** Coverage, duration, global explained variance (GEV), occurrence, and spatial correlation (SPC) for cases and controls with subgroup maps (left column) and combined maps (right column). * *p* < 0.05.

In order to test whether this finding was likely to be robust across microstate segmentation methods, we tested a single permutation using the analytic approach implemented in CARTOOL^16^ (for details, see^6^). We selected a permutation with a high degree of case-control spatial correlation between subgroup maps (all *r* > 0.95). In this specific analysis, we found that 66% of the comparisons were statistically significant using subgroup maps versus none when using shared maps. This suggests that our results are not specific to any one microstate segmentation procedure but rather a common feature of this kind of analysis.

Other factors may exacerbate or attenuate this source of bias, including i) choice of *K*, ii) sample size, iii) length of recordings and number of GFP peaks extracted, iv) number and layout of EEG channels, v) artifact/noise in the signals, vi) analytic choices in segmentation and backfitting, and vii) demographic/medical factors of participants. It is beyond the scope of this report to fully explore all such variables, especially as there may be complex effects: for example, a larger sample size will reduce the magnitude of chance group differences between estimated maps (thereby reducing bias) but on the other hand, a larger sample will also increase the statistical power to detect small differences in the derived metrics, whatever their source (thereby increasing bias). To address the choice of *K*, we repeated our primary set of simulations but using *K* = 6 segmentation; Figure 5B (top row) shows the combined maps, adopting the same microstate nomenclature as Figure 5 from ref. ^1^ (here the six maps correspond to A, B, C, C’, D & E). Median case-control spatial correlations were 0.92, 0.96, 0.98, 0.87, 0.83 and 0.96 respectively. We observed similar results as for the *K* = 4 simulations: inflated type I error rates despite generally high spatial correlations between case and control maps, even if restricting analyses to only maps with case-control spatial correlations *r* > 0.95. Across all replicates, one third of tests exhibited a type I error rate above 20% (maximum 53.4%). Experiment-wide, 46.2% of replicates were Bonferroni-significant, adjusting for 30 tests.

### Group differences in microstate topographies can confound analyses of derived metrics

The previous simulations demonstrated that group-specific segmentation can lead to spurious differences in microstate metrics, despite no true population-level differences in either microstate maps or metrics. In applications to real data however, two groups (or physiological states) may in fact differ in their characteristic microstate topographies. In this instance, analyses of group differences in microstate metrics that use only a single set of maps (whether derived from the entire dataset, a subset of it or an external template) may not be appropriate either, as underlying topographical group differences may be reflected as differences in microstate parameters such as coverage.

To illustrate this point, we conducted a proof-of-principle simulation to generate two groups that differed with respect to microstate map topographies but were otherwise identical. Specifically, we created a duplicate of the original *N* = 30 dataset which was identical except for having partially shuffled EEG channel labels, effectively “rotating” parts of each map (see Methods). As standard microstate segmentation is agnostic to channel location, these two datasets are identical through the lens of microstate analysis, except for different microstate topographies. We followed the procedure as above to estimate *K* = 4 subgroup specific and combined maps in each *N* = 30 dataset, both original and shuffled (Figure 4). As expected, the global variance explained was similar for subgroup segmentations (GEV = 0.72688 in both cases, with very small differences due to the random sampling of GFP peaks). In contrast, the combined (*N* = 60) segmentation yielded a slightly lower GEV of 0.711, reflecting the increased heterogeneity in the combined set. Maps were paired across groups based on the known, original labels, although empirically matching maps also led to the same pairings given high spatial correlations between original and shuffled maps (*r* = 0.93 for all 4 microstates). We then estimated the correlations in microstate metrics between the two groups: given we’ve paired equivalent maps, metrics should be perfectly correlated (*r* = 1.0) and show no group differences in means (allowing for trivial numerical differences due to random sampling of GFPs). This is indeed what we observed when using subgroup specific maps (Figures 4B & 4C). In contrast, all analyses based on a single set of maps (either a combined map, or based on one of the two groups) often showed correlations less than 1.0, and as low as 0.67 for some microstate metrics (Figure 4B), along with highly significant group differences in means (e.g. *p* ∼ 10^−8^ for B coverage from a comparison of *N* = 30/30). These results underscore the intuition that if the same underlying microstates have even quite modest group differences (i.e. here *r* = 0.93) in topographies, analyses that assume a universal set of maps can lead to spurious inferences about group differences in other microstate metrics. (Note that in this contrived example, using subgroup maps obligatorily yielded unbiased results only because the simulation procedure precluded stochastic sampling variation from inducing chance differences in microstate maps.)

**Figure 4.**
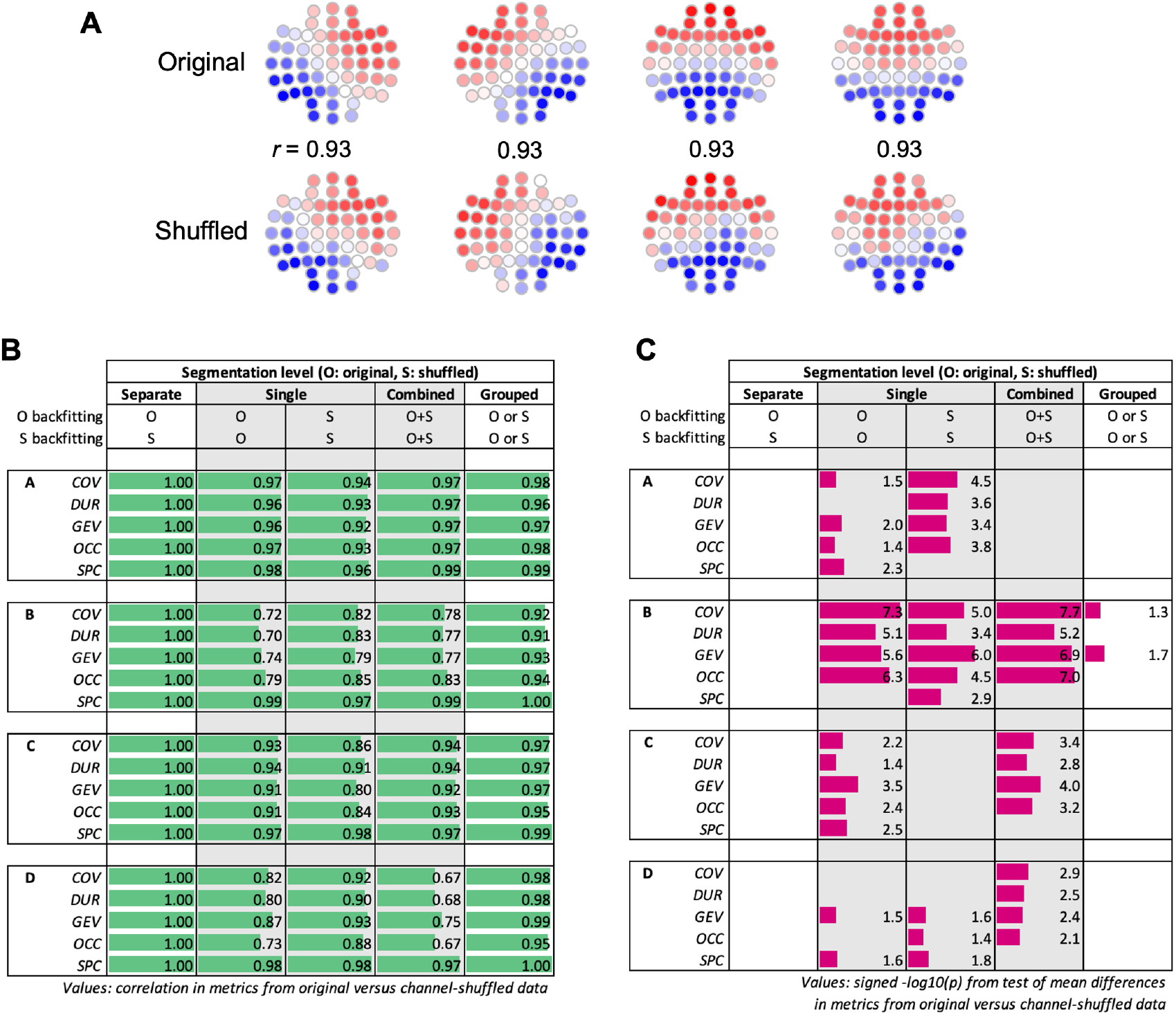
Impact of true group differences in microstate maps when using a combined/single set of maps. In a proof-of-principle single simulation based on the original *N* = 30 dataset, we created a second dataset which was identical except for having partially shuffled EEG channel labels. As the standard microstate segmentation procedure does not consider channel location, these two datasets are effectively identical through the lens of microstate analysis (except for the different microstate topographies). **A)** Maps from a *K* = 4 segmentation of the original (upper row) and channel-labeled shuffled datasets (lower) along with the spatial correlations between corresponding maps. **B)** Correlations between microstate metrics in the original (O) and shuffled (S) datasets (which should in principle be 1.0); using a single/combined set of maps leads to lower correlations. **C)** Unsigned -log10(*p*) significance values between metrics from the original and channel-shuffled dataset (showing *p*<0.05 results only). Single/combined maps show significant differences in microstate metrics (due to unmodelled differences in microstate maps rather than true differences in microstate dynamics).

### A grouped backfitting approach to alleviate bias in group comparisons

To test for group differences in derived microstate metrics whilst allowing for potential group differences in microstate topographies, a possible approach is to segment subgroup maps separately but then backfit all individuals to the total, grouped set (e.g *K* = 4 + 4 = 8), appropriately merging labels when calculating metrics such as coverage such that they represent *either* the case-derived state *A*, or the control-derived state *A*, for example. For a *K* = 4 segmentation, this would entail separately estimating maps in cases, e.g. {A_1_, B_1_, C_1_ and D_1_} and controls, e.g. {A_2_, B_2_, C_2_ and D_2_}, with labels assigned based on similarity to an external template. All individuals (both cases and controls) are backfit in the usual manner to the grouped set of eight maps {A_1_, A_2_, B_1_, B_2_, C_1_, C_2_, D_1_ and D_2_}. However, coverage, duration and occurrence metrics are estimated only for the common microstate, e.g. A, assuming that A_1_ and A_2_ are interchangeable, so that “coverage of A” implies “coverage of either A_1_ or A_2_”. Likewise, the mean duration of A is based on contiguous stretches of either A_1_ or A_2_. For meaningful interpretation, this assumes that A_1_ and A_2_ reflect topographically variable manifestations of the same underlying microstate, rather than two completely distinct microstates.

In our primary, original simulations (which did not induce topographical group differences, i.e. “same maps/same metrics”), we observed expected type I error rates for both *K* = 4 and *K* = 6 scenarios when using this approach (data not shown). When applied to the above “different maps/same metrics” simulation, spurious differences between groups were greatly attenuated, albeit not completely eliminated (Figure 4C). More work will be required to determine the properties of this approach in the general case.

### Application to real data: waking versus sleep microstates

Finally, we applied all approaches described above (separate subgroup maps, combined maps, grouped maps as well as single-group maps) to an analysis of real data, comparing the original *N* = 30 set of waking (quiet rest) recordings to a matched set of recordings obtained during N2 sleep on the same individuals (Figure 5). In contrast to the prior simulations conducted under the null hypothesis, here we expected actual differences in microstate topographies and/or dynamics between the two arousal states. As an initial *K* = 4 segmentation yielded qualitatively different sets of maps between wake and sleep (Figure 5A), we focused all analyses on a *K* = 6 segmentation which yielded more similar sets of maps (Figure 5B, using the A, B, C, C’, D & E nomenclature from ref. ^1^). Spatial correlations between wake and sleep maps were high (*r* > 0.9) except for microstate D (*r* = 0.79).

**Figure 5.**
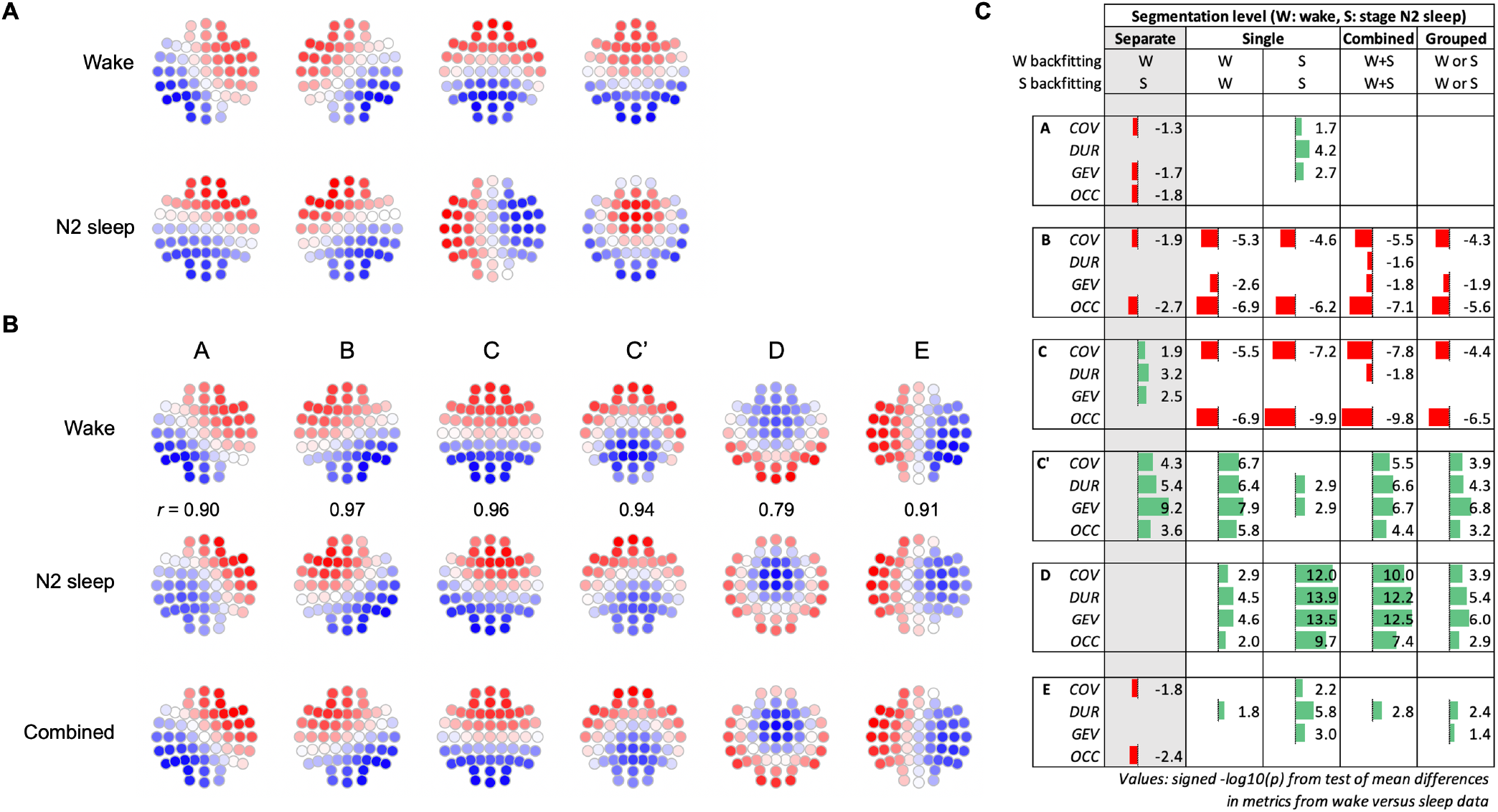
Application to real data comparing waking (quiet rest) and stage N2 sleep. A) *K* = 4 maps for wake and stage N2 sleep in those same individuals, yielding qualitatively different maps in this particular set of *N* = 30 healthy controls. B) *K* = 6 maps for wake (top row), stage N2 sleep (middle row) or from a segmentation on data combined across states (bottom row). Spatial correlations reflect the similarity of wake and sleep maps, which are generally high except for microstate D. C) Between group (wake vs sleep) analyses based on the K=6 segmentation; for nominally significant (*p* < 0.05) tests, values indicate the signed -log10(*p*) from a paired-sample *t*-test, with positive (green) values indicating higher values during sleep compared to wake. COV = coverage, DUR = duration, GEV = global variance explained, OCC = occurrence. Results for spatial correlations not shown as all methods showed similarly significant results for all microstates (higher values in sleep).

We performed backfitting and subsequent intra-individual tests for group differences in five ways (Figure 5C). The first column shows results using separate subgroup maps (backfitting wake recordings to wake-based maps, and sleep recordings to sleep-based maps), an approach we’ve shown is biased under the global null. In contrast, all four other approaches applied a common set of maps to all recordings. Focusing on combined and grouped maps, we observed a relatively high degree of consistency in results: broadly, reduced activity of microstates B and C during N2 sleep, and increased activity of C’, D and E.

In contrast, for microstates A, C, and E separate subgroup analyses yielded results that were significant but qualitatively divergent from combined/grouped analyses. For microstate D, which showed the strongest group differences for all other approaches, separate subgroup analyses yielded no significant differences. Consistent with the simulation results under the null, this suggests that biases arising from subgroup-specific segmentation can also attenuate true results or induce spurious results when the alternative hypothesis is true.

Analyses using single or combined maps likely confound differences in topography with differences in temporal metrics, which was most evident for microstate D (which showed the largest differences in topography between wake and sleep). In contrast, grouped analysis (last column) explicitly allows for group differences in microstate topography and so showed results that were qualitatively similar but less likely to be influenced by topographical differences. Thus, although grouped analysis tended to show reduced statistical significance, we suspect this is because they reflect a purer test of microstate dynamics.

## Conclusions

We show that, when microstate maps are estimated separately within subgroups, small differences in microstate topography — *arising purely by chance* — can nonetheless induce detectable differences in microstate parameters and lead to increased rates of spurious differences between groups. We suggest that group differences in microstate parameters be calculated using a single, common set of maps to eliminate this bias. Type I error rates should be appropriately controlled under the null hypothesis of no group differences in either microstate topographies or dynamics. A common set can be estimated from all the data (e.g. cases and controls), but could instead be derived from a subset (e.g. controls only) or based on pre-defined templates estimated in an external dataset. Illustrating the latter approach, Giordano et al.^10^ used a single set of microstate templates derived from a large dataset of healthy controls in a study of microstate parameters in patients with schizophrenia (albeit using templates based on only 19 channels, which recent work suggests may be inherently unreliable^17^).

On the other hand, when groups do in fact differ in microstate topographies, using a common set of maps may not be sufficient to control type I error rates in subsequent tests of metrics, either. We therefore introduced the “grouped maps” approach, to allow for between-group differences in map topographies whilst testing for differences in other metrics. Although this appears to be a promising approach, we nonetheless stress that if two groups show i) large differences in microstate topographies, and/or ii) large differences in the mean spatial correlation of samples to their assigned map for a given microstate, one should generally avoid making strong inferences about differences (or lack thereof) in other derived parameters.

In summary, using subgroup specific maps should be avoided as it is fundamentally anti-conservative under the global null. Using a common set of maps is preferred, although we’ve demonstrated the potential for conflating topographic group differences with other effects. Without carefully considering potential topographic differences, significant results based on a simple common set of maps only indicates that *some* aspect of microstate topography and/or dynamics differs between groups. Using a common set of all subgroup maps and pairing assumed-equivalent maps potentially provides a more robust general solution. We suggest that this and other models should be investigated to deal with biases in group analyses of microstates.

## Materials and Methods

### Participants and Procedure

The analyzed data was collected as part of the Global Research Initiative on Neurophysiology of Schizophrenia (GRINS), a research collaboration between the Broad Institute of Harvard and the Massachusetts Institute of Technology (USA), Harvard Medical School (USA), and Wuxi Mental Health Center (China). All study procedures were approved by the Institutional Review Board of Harvard TH Chan School of Public Health and the Institutional Review Board of the Wuxi Mental Health Center. Data collection procedures for this study were previously reported in ^18^. All participants were between 18 and 45 years old, had no history of serious neurologic or psychiatric illness, and were not taking medications. All participants provided informed consent before completing the study. For each participant, eight minutes of eyes-closed resting states EEG data was recorded using a 64-channel Ag/AgCl EasyCap system (Brain Products GmbH, Germany). After semi-automatic artifact rejection (including segments with excessive eye movement), the final recording durations were 4.5 minutes on average (ranging from 2 to 7 minutes). Data were recorded at 500 Hz with a left clavicle ground and FPz reference electrode. Electrode impedances were kept below 10 kΩ. For further details, please see ^18^.

### Microstate segmentation

Microstate segmentation and fitting was performed using Luna (http://zzz.bwh.harvard.edu/luna/). For microstate segmentation, based on 57 average-referenced and bandpass filtered (0.3 to 35 Hz) EEG channels, we calculated the global mean field power (GFP) across all electrodes. For each recording, we i) identified all GFP local maxima (“GFP peaks”), ii) rejected peaks that were of unusually high (>3 standard deviation (SD) units above the GFP peak mean) or low (>1 SD below) power, iii) as well as peaks with high spatial kurtosis (>1 SD above mean GFP kurtosis), as can result if one or two electrodes have high amplitude artifacts. From the remaining set, in keeping with past work^16^, we extracted 1000 time points for the subgroup map condition; we extracted 500 time points per recording for the combined map condition, thereby ensuring that all segmentation analyses were based on the same total number of data points (i.e. 30×500 = 15×1000 = 15,000). For segmentation only, we concatenated GFP peaks across individuals and applied a polarity-insensitive *k*-means clustering with number of distinct states *K* = 4, following refs^1,19^. In subsequent sensitivity analyses, we repeated this procedure setting *K* = 6 instead.

### Automated labelling and alignment of microstates

This resulted in four microstate maps for each of three conditions: a single combined set (*N* = 30), and for each null replicate, a “case” (*N* = 15) and “control” (*N* = 15) set, each arbitrarily ordered and labelled 1, 2, 3 and 4. To align similar maps across conditions, we identified pairings between each set of four maps (1-4) and an external, labelled set of maps (A-D) previously derived from a larger set of equivalent EEG recordings. Considering all possible pairings (e.g. 1®B, 2®A, 3®D & 4®C), the optimal set was selected to minimize the total distance Σ_*i*_(1-*r*_*ij*_)^*x*^ where *i*=1,2,3,4 is the observed map, *j*=A,B,C,D is the putative matched template for that map, and *r*_*ij*_ is the corresponding spatial correlation. By default, we set *x=2* (i.e. to place more weight on a single dissimilar pair of maps, versus multiple only slightly divergent pairs, when calculating distance) although similar results were obtained with *x*=1. Each observed map was matched with exactly one template map and vice versa. We tracked if a replicate had a “suboptimal” match with the template, such that a map was assigned to one canonical microstate (e.g. B) but in fact had a higher spatial correlation with a different canonical microstate in the template (e.g. C, implying that one of the three other maps had an even higher spatial correlation with C). On average, 4.6% of assignments exhibited this type of pattern (more so for B, less so for A and C). Finally, as well as matching all three sets (i.e. case, control and combined maps) against a labelled template, we also matched each case and control map against the combined (*N* = 30) map, with equivalent results in terms of type I errors (data not shown).

### Testing for group differences in microstate parameters

For the backfitting to derive per individual microstate parameters, we labeled each time point in the original recordings with the closest microstate map (again, polarity-insensitive). Putative microstates of short (<20 milliseconds) duration were replaced by the next most likely state until all states were at least 20 milliseconds. Microstate duration was defined as the average amount of time a given map persisted before transitioning to another microstate. Microstate coverage was the portion of the data that was mapped to each microstate. Microstate global explained variance (GEV) was the amount of variance in the data that was explained by that microstate. Microstate occurrence was how often a given microstate appeared. Spatial correlation was the average correlation of all assigned maps to the microstate. Group differences were assessed using an independent-sample *t*-test, after removing any outliers (+/- 2 SD units from the mean).

### Grouped backfitting

We added a grouped option to Luna’s (v0.28) microstate command (MS), which internally pairs maps based on uppercase and lowercase labels (e.g. A and a, B and b, etc). When comparing two groups, if {A_1_, B_1_, C_1_, D_1_} are the maps derived from one group (encoded A, B, C and D) and {A_2_, B_2_, C_2_, D_2_} is the equivalent set from the second group (encoded a, b, c and d), Luna would backfit to the *K* = 4+4 = 8 maps but generate ultimate metrics for only the four unique microstates, by collapsing A and a, etc, into a single label when assessing microstate duration, etc, such that coverage of “A” reflects being in either A_1_ or A_2_. Likewise, variance explained and mean spatial correlation metrics are based on sample points for which either A_1_ or A_2_ is the most likely assignment but using the appropriate map dissimilarity/spatial correlation for either A_1_ or A_2_, whichever is the more likely. In principle, not all microstates need to be observed in both subgroup maps and subgroup maps need not be based on the same value of *K*, e.g. {A_1_, B_1_, C_1_ and D_1_} and {A_2_, B_2_, C_2_, D_2_ and E_2_} subgroup maps would be represented as a *K* = 4 + 5 = 9 set, which implicitly forces microstate E to have identical topographies between groups.

### Shuffled microstate topographies

For the proof-of-principle simulation described in Figure 4, to generate two groups with correlated but distinct topographies, we generated a dataset with partially shuffled maps, by “rotating” each of these sets (i.e. “rings” on the scalp) of channel labels by one position for the second group: { FCZ, FC2, C2, CP2, CPZ, CP1, C1, FC1 }, { FZ, F2, F4, FC4, C4, CP4, P4, P2, PZ, P1, P3, CP3, C3, FC3, F3, F1 } and { AFZ, AF4, F6, FC6, C6, CP6, P6, PO4, POZ, PO3, P5, CP5, C5, FC5, F5, AF3 }. Note that while *any* permutation of channel labels would have resulted in qualitatively similar simulation results, we performed this constrained shuffle to generate pairs of maps that were still highly correlated (*r >* 0.9) between groups, and such that A_1_ was more similar to A_2_ than to B_2_, C_2_ or D_2_, etc, thereby mirroring a typical real data context in which topographically similar maps would be paired with each other.

### Testing for group differences in microstate topographies

Prior to generating random case/control groupings, we also estimated *K* = 4 microstate maps separately for each individual (Figure 1). To assess topographical within- and between-group similarities based on these individual-level maps, we computed i) a 30-by-30 distance matrix, comparing each individual’s map to all others (using distance as defined above), and ii) each individual’s distance to the external template set of *K* = 4 maps. Then, based on these distances, we estimated a number of metrics to quantify potential global differences in microstate topographies between groups, separately for each null replicate: i) mean case-to-case distance, ii) mean control-to-control distance, iii) the absolute difference between these two values, iv) the ratio between the mean distance between phenotypically concordant pairs versus the mean distance between all phenotypically discordant pairs, v) mean case-to-template distance, vi) mean control-to-template distance, and vii) the absolute difference between these last two values. For each replicate, we randomly permuted case/control labels *R* = 1,000 times to estimate the distributions of these seven statistics under the null hypothesis that differences in microstate topography are independent of case/control status. If *Q* is the number of times a permuted statistic was equal to or greater than the observed statistic, the associated empirical *p*-value was estimated as (*Q*+1)/(R+1). As group labels were generated under the null (i.e. independent of microstate topography), we expected these seven tests to follow the distribution expected under the null, i.e. to be unbiased and conform to nominal type I error rates.

## GRINS Consortium Members

<u>Clinical Research Team:</u> Jun Wang, Chenguang Jiang, Guanchen Gai, Kai Zou, Zhe Wang, Xiaoman Yu, Guoqiang Wang, Shuping Tan, Michael Murphy, Mei Hua Hall, Wei Zhu, Zhenhe Zhou

<u>Molecular Genetics:</u> Lu Shen, Shenying Qin. Hailiang Huang

<u>Electrophysiology data analyses:</u> Nataliia Kozhemiako, Lei A Wang, Yining Wang, Lin Zhou, Shen Li, Jun Wang, Robert Law, Minitrios Mylonas, Michael Murphy, Robert Stickgold, Dara Manoach, Mei-Hua Hall, Jen Q. Pan, Shaun M. Purcell

<u>Project management:</u> Zhenglin Guo, Sinead Chapman, Hailiang Huang, Jun Wang, Chenaugnag Jiang, Zhenhe Zhou, Jen Q. Pan

<u>Principle Investigators:</u> Mei Hua Hall, Hailiang Huang, Dara Manoach, Jen Q. Pan, Shaun M. Purcell, Zhenhe Zhou

